# Epigenetic and Transcriptional Dysregulation in CD4+ T cells of Patients with Atopic Dermatitis

**DOI:** 10.1101/2021.12.03.471059

**Authors:** Amy A. Eapen, Sreeja Parameswaran, Carmy Forney, Lee E. Edsall, Daniel Miller, Omer Donmez, Katelyn Dunn, Xiaoming Lu, Marissa Granitto, Hope Rowden, Adam Z. Magier, Mario Pujato, Xiaoting Chen, Kenneth Kaufman, David I Bernstein, Ashley L. Devonshire, Marc E. Rothenberg, Matthew T. Weirauch, Leah Kottyan

**Affiliations:** Division of Allergy and Immunology, Cincinnati Children’s Hospital Medical Center, Cincinnati, OH; Division of Allergy and Clinical Immunology, Henry Ford Health System, Detroit, MI; Center for Autoimmune Genomics and Etiology, Cincinnati Children’s Hospital Medical Center, Cincinnati, OH; Division of Immunology, Allergy, and Rheumatology, University of Cincinnati, College of Medicine, Cincinnati, OH; Divisions of Biomedical Informatics and Developmental Biology, Cincinnati Children’s Hospital Medical Center, Cincinnati, OH; Department of Pediatrics, University of Cincinnati College of Medicine, Cincinnati, OH; Cincinnati Veterans Administration, Cincinnati, OH

**Keywords:** atopic dermatitis, functional genomics, NFκB, T cells, gene regulation, disease genetics

## Abstract

Atopic dermatitis (AD) is one of the most common skin disorders in children. Disease etiology involves genetic and environmental factors, with the 29 independent AD risk loci enriched for risk allele-dependent gene expression in the skin and CD4+ T cell compartments. We investigated epigenetic mechanisms that may account for genetic susceptibility in CD4+ T cells. To understand gene regulatory activity differences in peripheral blood T cells in AD, we measured chromatin accessibility (ATAC-seq), NFKB1 binding (ChIP-seq), and gene expression (RNA-seq) in stimulated CD4+ T cells from subjects with active moderate-to-severe AD and age-matched, non-allergic controls. Open chromatin regions in stimulated CD4+ T cells were highly enriched for AD genetic risk variants, with almost half of AD risk loci overlapping with AD-dependent ATAC-seq peaks. AD-specific open chromatin regions were strongly enriched for NFκB DNA binding motifs. ChIP-seq identified hundreds of NFKB1-occupied genomic loci that were AD-specific or Control-specific. As expected, the AD-specific ChIP-seq peaks were strongly enriched for NFκB DNA binding motifs. Surprisingly, Control-specific NKFB1 ChIP-seq peaks were not enriched for NFKB1 motifs, instead containing motifs for other classes of human TFs, suggesting a mechanism involving altered indirect NFKB1 binding. Using DNA sequencing data, we identified 63 instances of genotype-dependent chromatin accessibility at 36 AD risk variants (30% of AD risk loci) that could lead to genotype-dependent expression at these loci. We propose that CD4+ T cells respond to stimulation in an AD-specific manner, resulting in disease and genotype-dependent chromatin accessibility involving NFKB binding.

**AUTHOR SUMMARY:** Stimulated CD4+ T cells from patients with atopic dermatitis have disease-dependent regulation of how gene expression is regulated. This regulation is disease dependent and the way the DNA is accessible and the transcription factor NFKB1 binds is enriched for genetic risk variants. Clinically, the CD4+ T cells in the peripheral blood of patients with AD respond to stimulation in a disease and genotype-dependent manner.

## INTRODUCTION

Atopic dermatitis (AD) is one of the most common skin disorders in children, affecting nearly 20% of children worldwide, and contributing to significant social and financial burden for patients and their families [1]. Although AD often presents in childhood, up to 80% of patients with AD have persistent disease into adulthood [2, 3]. Currently, patients with moderate-to-severe AD are treated with a “one-size-fits-all” approach, but recent investigations have revealed several different AD endotypes [4]. Both genetic and environmental factors are clearly implicated in the pathogenesis of AD [5], with genome-wide association studies identifying 29 independent AD risk loci [6, 7].

Immunologically, AD involves skin barrier defects and CD4+ T cells that localize to the skin, producing inflammatory cytokines and amplifying epidermal dysfunction [8]. This can lead to allergic sensitization through a disrupted skin barrier and, ultimately, to the development of other allergic diseases along the atopic march including allergic rhinitis, food allergy, and asthma [9]. Recent studies suggest that early and aggressive management of AD may prevent allergic sensitization and further progression of the atopic march [10–12].

AD genetic risk variants are enriched for genes with genotype-dependent expression (i.e. expression quantitative trait loci (eQLTs)) in skin as well as CD4+ T cells. This study focuses on CD4+ T cells based on the critical role these cells have in shaping the immune response in AD and other allergic diseases. Notably, in transcriptional studies of food allergy, the most robust disease specific expression in CD4+ T cells has been detected after stimulation of CD3/CD28 or antigen-loaded dendritic cells [13–15], two stimulatory pathways that activate NFκB [16, 17]. NFκB signaling has a well-established role in AD by controlling the transcription of inflammatory cytokines such as IL6 as well as adhesion molecules such as ICAM-1, contributing to the inflammation seen in the skin as well as the disruption in the skin barrier [18, 19]. An important role for NFκB in CD4+ T cells in AD was established in a mouse model of AD in which mice injected with CD4+ T cells with inhibited NFκB signaling showed marked improvement in AD-like skin lesions compared to those injected with CD4+ T cells with a control vector [20].

Herein we hypothesized that AD loci may be epigenetically regulated. In order to test this hypothesis, we first focused on measuring the chromatin accessibility, NFKB1 binding, and gene expression in stimulated CD4+ T cells from subjects with active moderate-to-severe AD, along with age and ancestry-matched healthy, non-allergic controls. We identified 34,216 regions of chromatin across the genome that are accessible in an AD-dependent manner. These regions are highly enriched for DNA sequence motifs recognized by NFκB transcription factors. We therefore performed ChIP-seq for NFKB1 in the AD and control individuals, identifying 20,322 genomic loci with AD-dependent NFKB1 occupancy. Whole genome sequencing of these individuals and application of our MARIO method to identify allelic activity [21] revealed 63 instances of genotype-dependent chromatin accessibility at 36 AD risk variants that might lead to the genotype-dependent gene expression at these loci. Collectively, our finding demonstrate that the pathoetiology of AD involves epigenetic changes in CD4+ T cells, especially via NFκB-mediated gene expression regulated mechanism.

## RESULTS

We created a set of 3,143 AD-associated genetic risk variants at 29 independent risk loci (**Supplemental Table 1** and see **Methods**). Application of our RELI method [21] to these variants using expression quantitative trait locus (eQTL) data obtained from Genotype-Tissue Expression GTeX [22] and Database of Immune Cell Expression, Expression quantitative trait loci and Epigenomics (DICE) [23] revealed strongest enrichment for CD4+ T cells, along with skin (sun exposed and sun unexposed) (**Table 1**). This analysis indicates that alteration of gene regulatory mechanisms in CD4+ T cells is likely an important factor underlying AD-associated genetic risk.

**Table 1.**
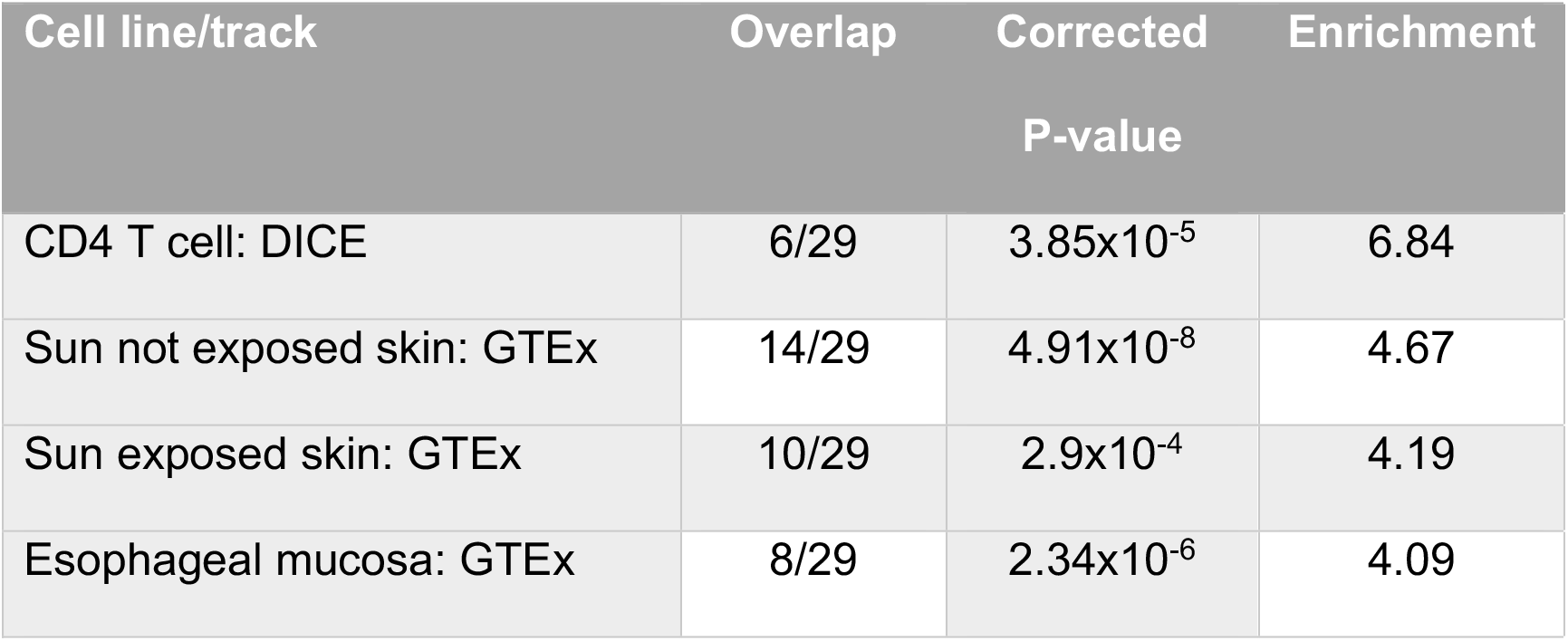
Enrichment of expression quantitative trait loci (eQTLs) at AD risk loci. Application of our RELI method [21] to 3,143 AD variants across 29 independent risk loci using expression quantitative trait locus (eQTL) data obtained from Genotype-Tissue Expression GTeX [22] and Database of Immune Cell Expression, Expression quantitative trait loci and Epigenomics (DICE) [23].

| Cell line/track | Overlap | Corrected P-value | Enrichment |
| --- | --- | --- | --- |
| CD4 T cell: DICE | 6/29 | $3.85 \times 10^{-5}$ | 6.84 |
| Sun not exposed skin: GTEx | 14/29 | $4.91 \times 10^{-8}$ | 4.67 |
| Sun exposed skin: GTEx | 10/29 | $2.9 \times 10^{-4}$ | 4.19 |
| Esophageal mucosa: GTEx | 8/29 | $2.34 \times 10^{-6}$ | 4.09 |

We recruited six moderate-to-severe AD patients (average EASI score of 30) and six age and ancestry-matched controls without known deleterious mutations in the *Fillagrin* gene. Demographics are indicated in **Table 2**. Adults with persistent AD had childhood onset of the disease. The mean total IgE among AD subjects (180.8 kU/L) was higher than among controls (61.7 kU/L). Peripheral blood was obtained from each subject and CD4+ T cells were isolated and stimulated for 45 hours with anti-CD3/CD28 beads (**Figure 1**).

**Figure 1:**
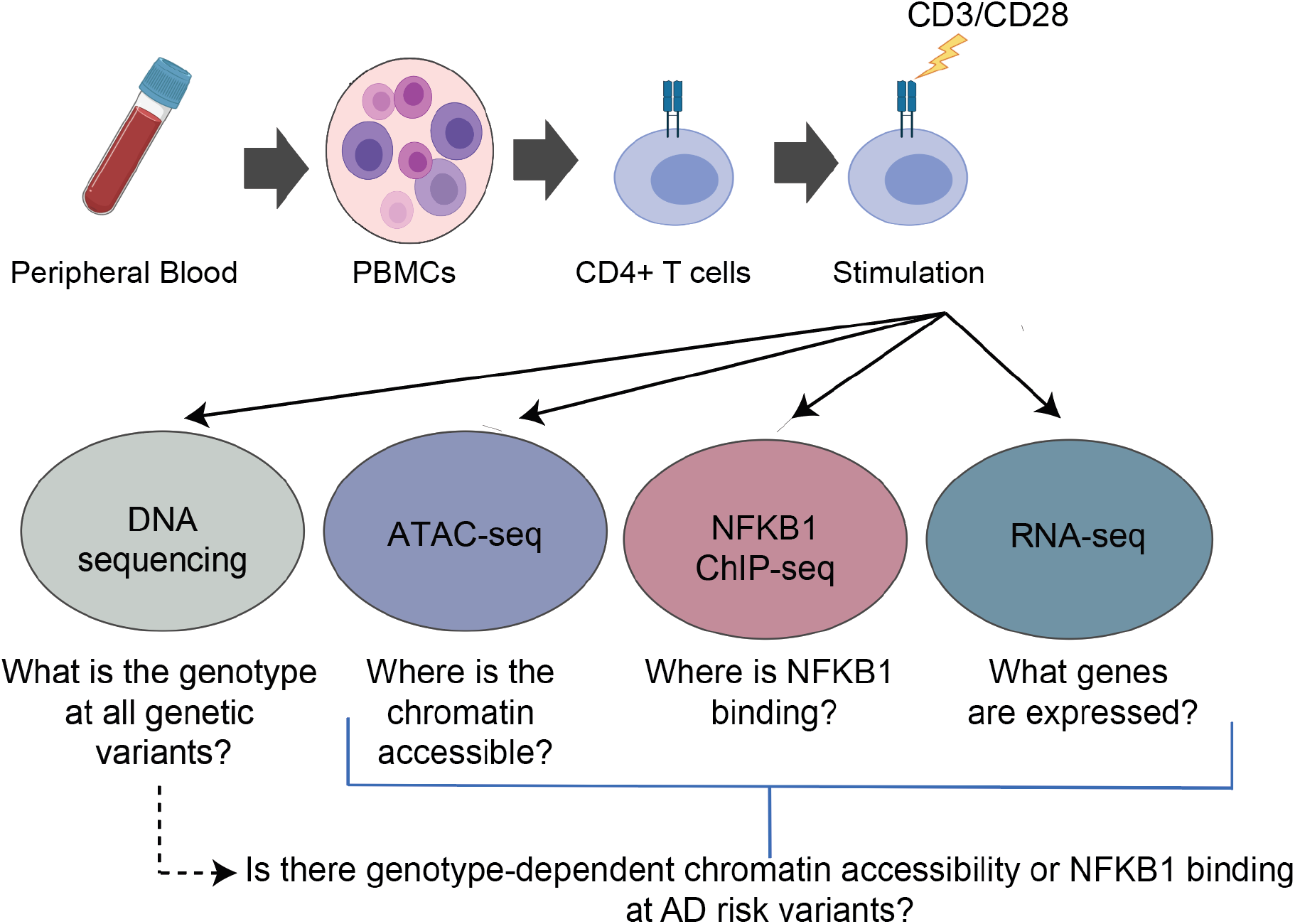
Study Design.

**Table 2.** Demographics on 6 age-matched AD cases and controls.

|  | Age (years) | Gender (% male) | EASI (0-72) | IGA (1-4) | Age of onset (years) | Asthma | Allergic rhinitis | Food allergy | Total IgE | Number of sensitizations |
| --- | --- | --- | --- | --- | --- | --- | --- | --- | --- | --- |
| Children with active AD (n=2) | 10 (8-12) | 1/2 (50%) | 31.9 (31.3-32.4) | 3.5 (3-4) | 4.5 (2-7) | 0 | 1/2 (50%) | 0 | 16.5 (6-27) | 3.5 (1-6) |
| Children without AD (n=2) | 13.5 (10-17) | 1/3 (33%) | n/a | n/a | n/a | n/a | n/a | n/a | 150.5 (23-278) | n/a |
| Adults with active AD (n=4) | 44.2 (25-67) | 1/4 (25%) | 29.2 (17.1-50.9) | 3.3 (3-4) | 4 (0.33-10) | 3/4 (75%) | 5/5 (100%) | 3/4 (75%) | 290.3 (18-648) | 5.3 (4-8) |
| Adults without AD (n=4) | 35.3 (18-53) | 0/4 (0%) | n/a | n/a | n/a | n/a | n/a | n/a | 17.2 (8-36) | n/a |

### Global mapping of the chromatin accessibility landscape in AD CD4+ T cells

We performed assay for Transposase-Accessible Chromatin followed by sequencing (ATAC-seq) to identify genome-wide chromatin accessibility. The data obtained were of high quality, with an average of almost 70,000 peaks per dataset, an average Fraction of Reads Inside of Peaks (FRiP) score of 0.32, and an average transcription start site (TSS) enrichment score of 20.5 **(Supplemental Table 2)**. Pairwise comparisons of each dataset identified strong agreement between subjects within cases and controls **(Supplemental Figure 1 A and B)**.

We assessed the the overlap of chromatin accessibility data with AD genetic risk variants using the RELI method [21]. Seven of the twelve ATAC-seq datasets were significantly enriched for AD risk loci with 7-14 overlapped risk loci for each subject (RELI p_corrected_: 0.01-1.6×10^−3^) (**Supplemental Table 3)**.

In a pairwise assessments performed using MAnorm [24], most ATAC-seq peaks were shared between AD patients and demographically matched controls (75.0-88.4%). The remaining peaks were either stronger in AD (AD-specific) or in Control (Control-specific) (representative analysis **Figure 2 A**, full results in **Supplemental Figure 2)**. We identified 34,216 regions of chromatin across the genome that were accessible in an AD-dependent manner, yielding 409 AD-specific and 398 Control-specific peaks that are present in three or more pairs (**Supplemental Figure 3**). We defined these ATAC-seq peaks that were AD-specific or control-specific in three or more subject pairs as “consistently AD-specific” and “consistently control-specific” peaks, respectively. Consistently AD-specific ATAC-seq peaks overlapped AD-associated genetic risk variants at 13 of the 29 AD risk loci (2.0-fold enrichment, p_corrected_=0.015) (**Supplemental Table 3)**. These results indicate that chromatin is accessible in a disease-specific manner in CD4+ T cells at almost half of the AD risk loci.

**Figure 2:**
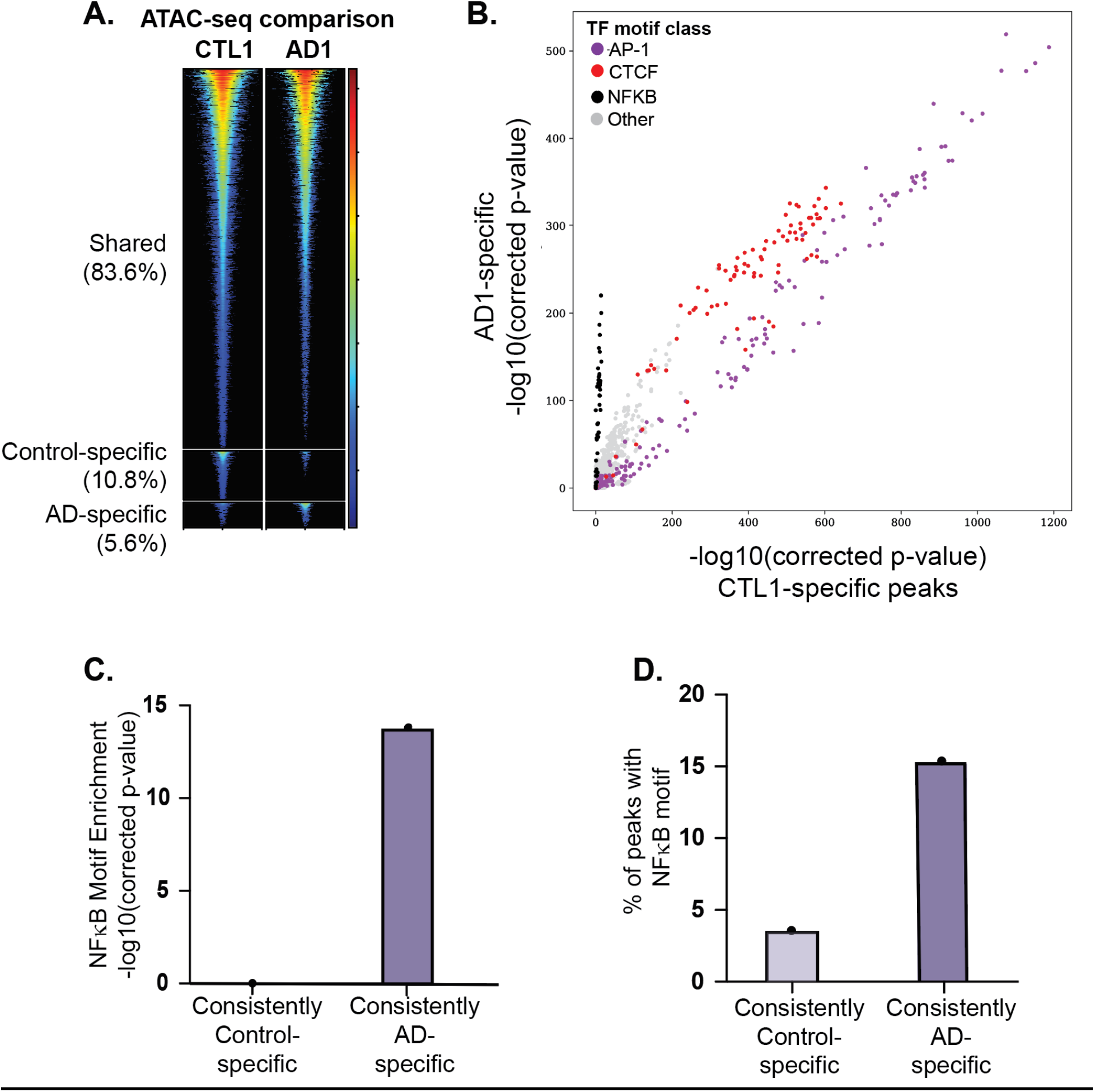
Differential chromatin accessibility and TF motif enrichment in AD subjects versus matched controls. ATAC-seq peaks were identified for all cases and controls and compared for all subject pairs. A. Differential chromatin accessibility analysis. For each matched pair of subjects, we identified shared peaks, Control-specific peaks, and AD-specific peaks (see Methods). A representative subject pair is shown in A. Each row represents a single genomic locus where an ATAC-seq peak was identified in either the AD or control subject. The center of each row corresponds to the center of the ATAC-seq peak. Heatmap colors indicate the normalized ATAC-seq read count within the AD1 subject (right) or CTL1 subject (left) – see key at right. B. Differential TF motif enrichment analysis. Comparison of TF motif enrichment results within a representative AD-specific and control-specific matched subject pair. Each dot represents the enrichment of a particular motif (corrected negative log 10 p-value). Select motif families are color-coded (see key at upper left). C and D. NFKB motif enrichment comparison between consistently AD-specific and consistently control-specific ATAC-seq peaks. “Consistently specific” peaks were defined as those peaks that were AD-or control-specific in at least three cases or controls, respectively. Results are shown for representative Cis-BP NFKB motif M05887_2.00. Full results are provided in **Supplemental Table 5**.

**Figure 3:**
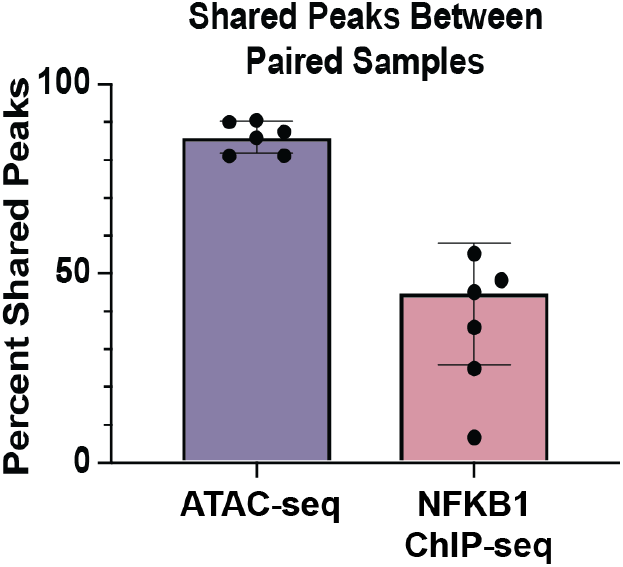
Shared peaks in ATAC-seq and ChIP-seq experiments between paired subjects. Percentage of shared peaks between paired samples in ATAC-seq compared to NFKB1 ChIP-seq experiments.

To identify potential transcription factors (TFs) whose binding might be affected by differential chromatin accessibility, we performed TF binding site motif enrichment analysis on AD- and control-specific ATAC-seq peaks. These analyses revealed that NFκB DNA binding motifs were more strongly enriched in the AD-specific vs control-specific ATAC-seq peaks in five of the six matched pairs, with the remaining pair showing equivalent enrichment for NFκB **(Figure 2B, Supplemental Figure 2, Supplemental Table 4)**. Motif enrichment analysis in consistently AD-specific or consistently control-specific peaks confirmed that NFκB binding motifs were highly enriched in an AD-specific manner, with ~15% of consistently AD-specific peaks containing predicted NFKB binding sites (P<10^−13^), and only ~3% of consistently control-specific peaks (P=1) (**Figure 2 C–D, and Supplemental Table 5**). These data highlight the potential for more robust direct binding of NFKB to the genome in activated CD4+ T cells in AD cases compared to matched controls.

### NFKB1 binds in an AD specific manner at hundreds of genomic loci in CD4+ T cells

We performed NFKB1 (p50) chromatin immunoprecipitation (ChIP-seq) experiments to measure NFκB binding to the genome of stimulated CD4+ T cells, obtaining an average of ~11,000 peaks per subject, with an average FRiP score of 0.012 **(Supplemental Table 2)**. All ChIP-seq peak datasets had highly significant overlap with a previously published CD4+ T cell NFKB1 ChIP-seq dataset (GSE126505) (**Supplemental Table 6**). As expected, the NFκB DNA binding motif was highly enriched in each of our NFκB ChIP-seq datasets (Cis-BP NFκB motif M05887_2.00 enrichment: 10^−4158^ < P < 10^−123^ **(Supplemental Table 7)**). We identified 20,322 genomic loci with AD-dependent NFKB1 occupancy. AD-specific NFKB1 ChIP-seq peaks were enriched for overlap with AD-specific ATAC-seq peaks in all six pairs (between 5.9 and 38.1-fold enrichment, 3.00×10^−25^ < p < 3.20×10^−203^) (**Supplemental Table 8**). There was substantially more variability between subject matched pairs in the NFKB1 ChIP-seq experiments compared to the ATAC-seq experiments, with a median of 51.5% shared NFKB1 peaks vs. a median of 91.9% shared ATAC-seq peaks (**Figure 3**). These results indicate substantially more differential NFKB1 binding than chromatin accessibility in AD subjects compared to matched controls.

We next sought to identify AD- and control-specific NFKB1 binding events by performing a pairwise assessment of NFKB1 peaks in cases vs controls (see Methods). This procedure identified shared, control-specific, and AD-specific NFKB1 peaks. An exemplary pair shown is in **Figure 4A**, with all pairs shown in **Supplemental Figure 4**. Strikingly, NFκB binding sites were more strongly enriched in the AD-specific NFKB1 ChIP-seq peaks compared to the matched control in five of the pairs **(Supplemental Figure 4** and **Supplemental Table 7**, example pair shown in **Figure 4B)**. We defined those NFKB1 peaks that were AD-specific or control-specific in three or more subject pairs as “consistently AD-specific” and “consistently control-specific” peaks, respectively (**Supplemental Figure 3 C-D**). In total, we identified 143 and 80 AD-specific and control-specific NFKB1 ChIP-seq peaks, respectively. Motif enrichment analysis revealed that NFKB binding motifs were also the most highly enriched motif class within consistently AD-specific NFKB1 peaks (**Figure 4 C–D** and **Supplemental Table 9**). In strong contrast, consistently control-specific peaks were not enriched for NFKB motifs, instead enriching for a wide range of motif classes (**Supplemental Figure 5 A-B** and **Supplemental Table 9**). Collectively, these results indicate that AD-specific NFKB ChIP-seq peaks strongly enrich for NFKB1 motifs, while control-specific NFKB peaks surprisingly do not.

**Figure 4:**
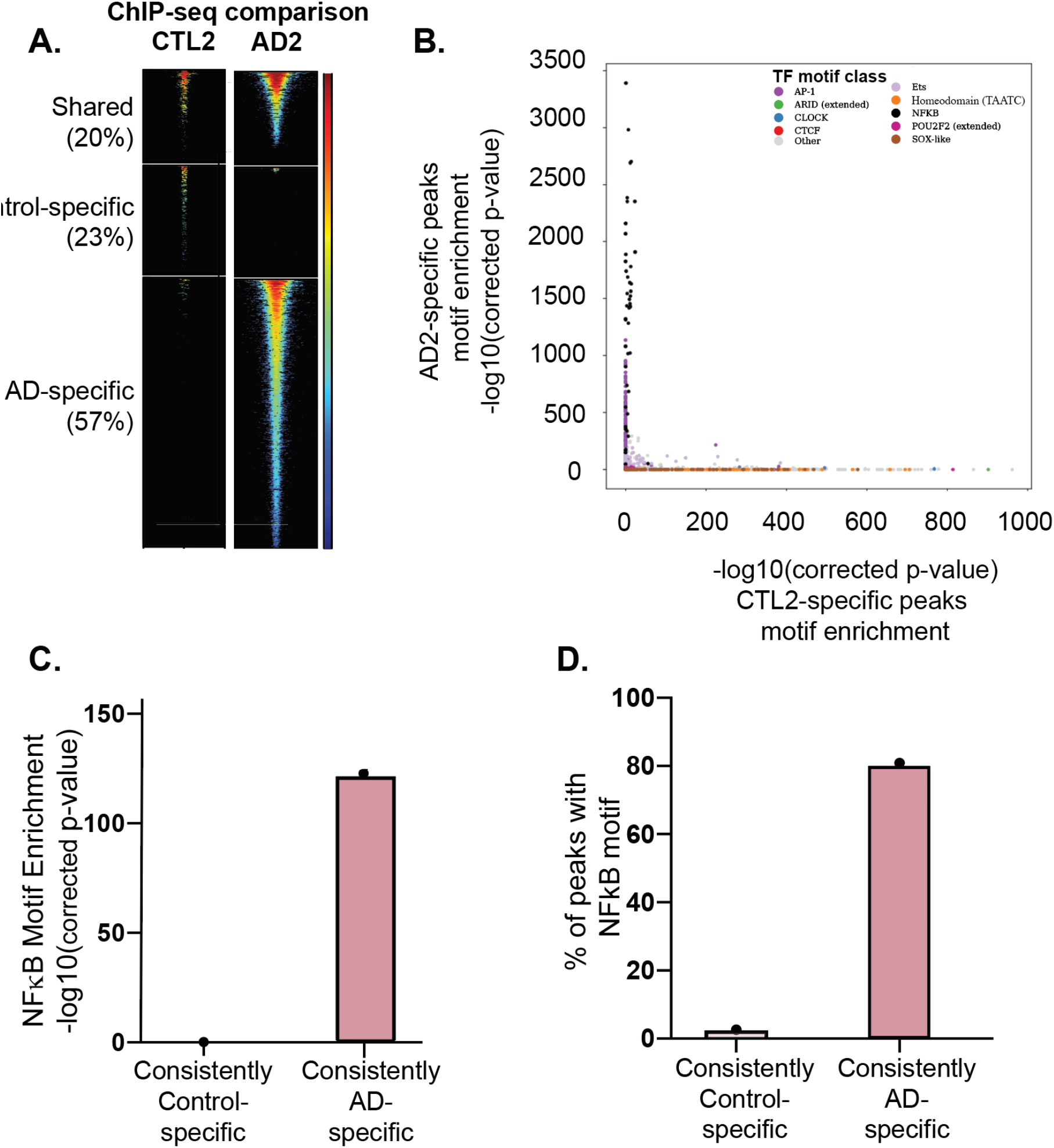
Differential NFKB1 binding and TF motif enrichment in AD subjects versus matched controls. NFKB1 ChIP-seq peaks were identified for all cases and controls and compared between matched subject pairs. A. Differential NFKB1 binding analysis. For each matched pair of subjects, we identified shared peaks, Control-specific peaks, and AD-specific peaks (see Methods). A representative subject pair is shown in A. Each row represents a single genomic locus where an NFKB1 ChIP-seq peak was identified in either the AD or control subject. The center of each row corresponds to the center of the ChIP-seq peak. Heatmap colors indicate the normalized ChIP-seq read count within the AD2 subject (right) or CTL2 subject (left) – see key at right. B. Differential TF motif enrichment analysis. Comparison of TF motif enrichment results within a representative AD-specific and control-specific matched subject pair. Each dot represents the enrichment of a particular motif (corrected negative log 10 p-value). Select motif families are color-coded (see key at upper right). C and D. NFKB motif enrichment comparison between consistently AD-specific and consistently control-specific NFKB1 ChIP-seq peaks. “Consistently specific” peaks were defined as those peaks that were AD- or control-specific in at least three cases or controls, respectively. Results are shown for representative Cis-BP NFκB motif M05887_2.00. Full results are provided in **Supplemental Tables 7** and **9**.

**Figure 5.**
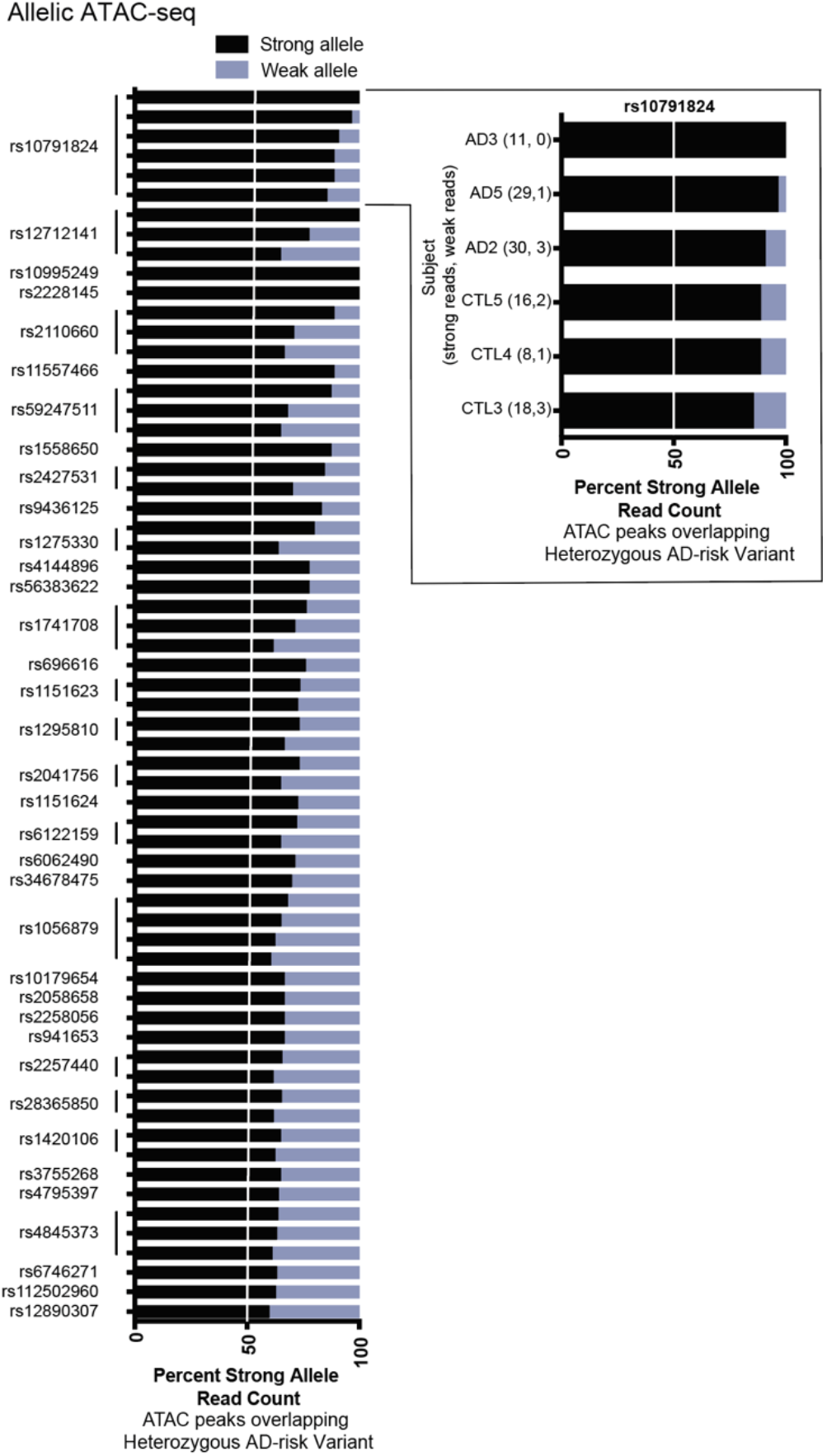
Allele-dependent chromatin accessibility at AD risk loci. A. AD-associated genetic risk variants with allele-dependent ATAC-seq peaks in CD4+ T cells. Each variant is heterozygous and located within an ATAC-seq peak in the indicated individual, enabling MARIO analysis to identify allele-dependent behavior. Full data are presented in **Supplemental Table 12**. All results shown have MARIO ARS values > 0.4 and are hence allele-dependent. In the cutout, the participant identifier and reads under the ATAC-seq peak overlapping rs10791824 mapping to the strong and weak bases are provided.

### Comparison across data types confirms strong concordance between RNA-seq, ATAC-seq, and NFKB1 ChIP-seq

We next measured gene expression levels in CD3/CD28-stimulated CD4+ T cells from each subject in this study, with the goal of integrating these data with chromatin accessibility and NFKB1 binding. In a case-control pairwise analysis, 15 genes were expressed at least 1.5-fold higher in the stimulated CD4+ T cells from the patient with AD compared to the matched control, and 16 genes were expressed 1.5-fold lower in the case compared to the matched control (**Supplemental Table 10**). These 31 genes were enriched for AD-related processes such as the “regulation of immune system processes”, “lymphocyte activation”, and “cytokine-mediated signaling pathway” GO biological processes as well as “cytokine receptor binding”, “nitric oxide synthase binding”, and “RNA-polymerase II-specific DNA-binding transcription factor binding” GO molecular functions (**Supplemental Figure 6**).

The 100 kB region of DNA around AD-specific gene sets widely overlapped (94.7-100%) the ATAC-seq peaks in the six subjects with AD (**Supplemental Table 8**). There was substantial overlap (26.3-68.4%) between the 100 kb region of DNA around AD-specific gene sets with the AD-specific ATAC-peaks, indicating that possible enhancers proximal to the AD-specific genes were accessible for transcription in an AD-specific manner. Similarly, the 100 kb region of DNA around NFKB1 ChIP-seq peaks overlapped the transcriptional start site of 47-95% of the AD-specific genes (**Supplemental Table 8**). In five of the six pairs, AD-specific NFKB1 ChIP-seq peaks overlapped a large proportion of the AD-specific genes (42.1-73.4%) (**Supplemental Table 8**). Collectively, these data indicate strong agreement between AD- and control-specific gene expression, chromatin accessibility, and NFKB1 binding.

### Allele-dependent chromatin accessibility at AD risk loci

Increasing evidence points to an important role for allele-dependent gene regulatory mechanisms in many diseases [21, 25–27]. To identify such events, we performed whole genome sequencing of all subjects to identify the alleles present at AD genetic risk variants (**Supplemental Table 11**). We integrated these data with the functional genomics data produced in this study using the Measurement of Allelic Ratios Informatics Operator (MARIO) method, which measures the allele-dependence of sequencing reads at genetic variants that are heterozygous [21]. Collectively, there were an average of 2.3 (0-5) heterozygous loci in AD cases and 2.6 (range 0-6) in controls (**Supplemental Table 11**), providing 124 total opportunities to discover allele-dependent ATAC-seq or NFKB1 peaks at AD genetic risk variants.

Sixty AD-associated variants are located within an ATAC-seq peak in at least one subject and also heterozygous in that subject. Strikingly, 36 of these 60 (60%) variants produced allele-dependent ATAC-seq peaks (**Supplemental Table 12, Figure 5**). Collectively, the AD risk variants with allelic chromatin accessibility were found at nine independent risk loci (31% of AD risk loci). At 16 of the AD risk variants that were heterozygous and overlapped ATAC-seq peaks, we measured allelic imbalance across multiple subjects. For example, at rs10791824 near the *OVOL1* gene, we measured a strong preference for the A allele across six individuals, with 85-100% of reads for all subjects having an A (total of 112 vs 10, A vs T reads). 28 of the AD risk variants with allelic ATAC were found to be eQTLs in stimulated CD4+ T cells based on DICE as curated by the eQTL catalogue [28] (**Supplemental Table 13**); however, these associations of allelic expression were not robust to multiple testing correction after accounting for all of the eQTL measurements in that study (i.e. across many cell types with and without stimulation).

In contrast to the large amount of observed allelic ATAC-seq peaks, only 6 unique AD-associated variants were located within at least one NFKB1 ChIP-seq peak and also heterozygous in any of the 12 subjects, and none of these demonstrated genotype-dependent activity. Future studies examining larger cohorts will be better powered to potentially identify allelic NFKB1 binding activity.

## DISCUSSION

Altogether, our data support a model in which stimulated peripheral blood CD4+ T cells from patients with active AD have extensive differential chromatin accessibility and NFKB1 binding relative to matched controls. We identify genotype-dependent chromatin accessibility at 36 AD genetic risk variants. Collectively, this study highlights plausible genetic risk mechanisms for AD through disease-specific epigenetic factors that are enriched at AD genetic risk loci. Current databases of eQTLs are not specific to participants with AD or other allergic diseases and they do not contain sufficient numbers of participants to have the statistical power to robustly identify moderately sized eQTLs. It will be important for the field to continue to curate control and AD-specific eQTL datasets. It will also be important to perform molecular studies assessing genotype-dependent regulatory activity at AD risk variants in the context of CD4+ T cells to definitively identify allelic transcriptional mechanisms at these loci.

In this study, we use a strong stimulation (CD3/CD28 crosslinking) to model immune activation in patients. It is possible that differing levels of stimulation will reveal additional disease specific differences in future studies. We used CD4+ T cells in our functional genomic assays to support quantitative assessment of disease-specific and genotype-dependent mechanisms. The disease and genotype-dependent effects observed in this study can be refined and validated in very specific immune cell subsets as the technology and analytical framework for quantitative comparisons at a single cell level continue to mature.

Differences in the subtypes of CD4+ T cells in the peripheral blood of patients with moderate to severe AD might explain some of the differential functional genomic effects found in this study. Recent investigations have revealed that patients with AD have an expansion of the T helper type 9 (Th9) subset of CD4+ T cells, and increased frequency of circulating CD25hiFoxp3+ T cells, compared to controls [29–31]. Skin-homing Th22 T cells are also increased in patients with AD across ages [30]. Additionally, the T cell profile naturally changes with age, and these changes are different in patients with AD [32]. Natural aging accounts for both quantitative and qualitative changes in the CD4+ T cell compartment [33]. Although our study did not differentiate different CD4+ T cell subtypes, our case-vs-control comparisons were all performed between demographically matched controls. Future studies could investigate the role of these different subtypes by age and the prevalence of genotype-dependent transcriptional dysregulation in AD patients compared to controls.

Current treatment of AD includes topical steroids, aggressive moisturization, anti-inflammatory non-steroidal agents (e.g. anti-type 2 immunity biological agents and JAK inhibitors), and in severe cases, systemic immunosuppression. There is a recognized need for newer therapeutics to address the complexity of the disease and for personalized medicine for subtypes/phenotypes differing by age, disease chronicity, and underlying molecular mechanisms [4, 34]. The results of this study support a continued focus on therapeutics aimed at the inhibition of NFκB-signaling. Numerous studies in mice and humans provide further rationale for focusing on this mechanism in AD [35–37].

In conclusion, the results of this study support a model in which stimulated CD4+ T cells from patients with AD have disease and allele-dependent differences in chromatin accessibility, and disease-dependent differences in NFKB1 binding. Based on the broad genotype-dependent chromatin accessibility at AD risk variants in stimulated CD4+ T cells, our data support allelic transcriptional regulation as an important epigenetic mechanism mediating disease risk.

## METHODS

### Collection of AD-associated genetic risk variants

122 genetic variants reaching genome-wide significance in a GWAS of AD were identified from the Genome Wide Association (GWAS) Catalogue (https://www.ebi.ac.uk/gwas/) [38] and a genetic association study on the Illumina ImmunoChiP [39]. Independent risk loci were identified through linkage disequilibrium pruning (*r*^*2*^ < 0.2) to identify a total of 29 genetic risk loci. We identified 3,143 AD risk variants across these 29 loci by accounting for linkage disequilibrium (*r*^2^ > 0.8) based on 1000 Genomes Data [40] in the ancestry(ies) of the initial genetic association using PLINK(v1.90b) [41] (**Supplemental Table 1**).

### Patient recruitment

Patients with moderate-to-severe AD were recruited from Cincinnati Children’s Hospital Allergy clinics, the Bernstein Allergy Group, and the Bernstein Clinical Research Center for this IRB approved study. Matched healthy controls were recruited by advertisement. To reduce heterogeneity, the study inclusion criteria for the subjects with AD were: 1) Presence of atopy established by positive aeroallergen skin prick testing and/or elevated serum total IgE; and 2) Moderate to severe AD defined as an Eczema Area and Severity Index (EASI) score ≥17 and Investigator Global Assessment (IGA) score ≥3 [42]. These AD severity tools have been previously validated for AD severity scoring [42, 43]. Control subjects were included if they had no history of atopic disease with negative aeroallergen skin prick testing as performed at their enrollment visit. Exclusion criteria included history of being on any biologic therapy, oral steroids or immunosuppressive medications in the past 6 months, due to their effect on T cells and the transcriptome.

### CD4+ T cell stimulation

After meeting inclusion and exclusion criteria, we isolated peripheral blood mononuclear cells (PBMCs) using Ficoll-Paque (GE Healthcare) density gradient separation from AD and control individuals. 51.5 million PBMCs ± 20.1 million were isolated from each subject. CD4+ T cells were then isolated using magnetic column separation (Miltenyi Biotec, CD4+ T cell isolation kit, human); 17.7 million CD4+ T cells ± 7.2 million were isolated from each subject. To activate NFκB, we stimulated these CD4+ T cells with CD3/CD28 crosslinking for 45 hours (Gibco, Dynabeads Human T-Activator CD3/CD28). We then performed ChIP-seq (2 million stimulated CD4+ T cells), ATAC-seq (50,000 stimulated CD4+ T cells), and RNA-seq assays (2.5 million stimulated CD4+ T cells), see below. Whole genome sequencing identified genetic variants.

### Assay for Transposase-Accessible Chromatin sequencing (ATAC-seq)

Briefly, transposase Tn5 with adapter sequences was used to cut accessible DNA [44]. These accessible DNA with adaptor sequences were isolated, and libraries were prepared from 50,000 stimulated CD4+ T cells using the OMNI ATAC protocol [45]. The libraries were sequenced at 150 bases per end on an Illumina NovaSeq 6000 at the Cincinnati Children’s Hospital Medical Center (CCHMC) DNA Sequencing and Genotyping Core Facility. The quality of the sequencing reads was verified using FastQC (version: 0.11.2) (http://www.bioinformatics.babraham.ac.uk/projects/fastqc) and adapter sequences were removed using cutadapt (trimgalore version: 0.4.2). ATAC-seq reads were aligned to the human genome (hg19) using Bowtie2 [46]. Aligned reads were then sorted using samtools (version 1.8) [47] and duplicate reads were removed using picard (version 1.89) (https://broadinstitute.github.io/picard/). Peaks were called using MACS2 (macs2 callpeak -g hs -q 0.01) [48]. ENCODE blacklist regions (https://github.com/Boyle-Lab/Blacklist/tree/master/lists/hg19-blacklist.v2.bed.gz) [49] were removed. Differential chromatin accessibility was calculated using the MAnorm program [24] with thresholds of fold change greater than 1.5 and p-value less than 0.05.

### Chromatin immunoprecipitation sequencing (ChIP-seq)

CD4+ T cells from subjects were crosslinked and nuclei were sonicated. Cells were incubated in crosslinking solution (1% formaldehyde, 5 mM HEPES [pH 8.0], 10 mM sodium chloride, 0.1 mM EDTA, and 0.05 mM EGTA in RPMI culture medium with 10% FBS) and placed on a tube rotator at room temperature for 10 minutes. To stop the crosslinking, glycine was added to a final concentration of 0.125 M and tubes were rotated at room temperature for 5 minutes. Cells were washed twice with ice-cold PBS, resuspended in lysis buffer 1 (50 mM HEPES [pH 8.0], 140 mM NaCl, 1 mM EDTA, 10% glycerol, 0.25% Triton X-100, and 0.5% NP-40), and incubated for 10 min on ice. Nuclei were harvested after centrifugation at 5,000 rpm for 10 min, resuspended in lysis buffer 2 (10 mM Tris-HCl [pH 8.0], 1 mM EDTA, 200 mM NaCl, and 0.5 mM EGTA), and incubated at room temperature for 10 minutes. Protease and phosphatase inhibitors were added to both lysis buffers. Nuclei were then resuspended in the sonication buffer (10 mM Tris [pH 8.0], 1 mM EDTA, and 0.1% SDS). A S220 focused ultrasonicator (COVARIS) was used to shear chromatin (150-to 500-bp fragments) with 10% duty cycle, 175 peak power, and 200 bursts per cycle for 7 minutes. A portion of the sonicated chromatin was run on an agarose gel to verify fragment sizes. Sheared chromatin was precleared with 10 μl Protein G Dynabeads (Life Technologies) at 4 °C for 1 hour.

Immunoprecipitation of NFKB1-chromatin complexes was performed with an SX-8X IP-STAR compact automated system (Diagenode). Beads conjugated to antibodies against NFKB1 (Cell Signaling (D7H5M) Rabbit mAb #12540) were incubated with precleared chromatin at 4°C for 8 hours. The beads were then washed sequentially with buffer 1 (50 mM Tris-HCl [pH 7.5], 150 mM NaCl, 1 mM EDTA, 0.1% SDS, 0.1% NaDOC, and 1% Triton X-100), buffer 2 (50 mM Tris-HCl [pH 7.5], 250 mM NaCl, 1 mM EDTA, 0.1% SDS, 0.1% NaDOC, and 1% Triton X-100), buffer 3 (2 mM EDTA, 50 mM Tris-HCl [pH 7.5] and 0.2% Sarkosyl Sodium Salt), and buffer 4 (10 mM Tris-HCl [pH 7.5], 1 mM EDTA, and 0.2% Triton X-100). Finally, the beads were resuspended in 10 mM Tris-HCl (pH 7.5) and used to prepare libraries via ChIPmentation [50].

The ChIP-seq libraries were sequenced as single end, 100 bases, on an Illumina NovaSeq 6000 at the CCHMC DNA Sequencing and Genotyping Core Facility. The reads were processed and analyzed as described above for ATAC-seq. We also used publicly available NFKB1 ChIP-seq datasets (GSE126505), which were processed using the same analytical pipeline.

### RNA-sequencing (RNA-seq)

Total RNA was extracted using the mirVANA Isolation Kit (Ambion) from stimulated CD4+ T cells of controls and AD subjects 45 hours post stimulation. RNA-seq libraries were sequenced as paired end, 150 bases. FastQC and cutadapt were used to verify read quality and remove adapters as above. RNA-seq reads were aligned to the hg19 (GrCh37) genome build (NCBI) using Spliced Transcripts Alignment to a Reference (STAR, version: 2.5.2a) [51]. The program featureCounts (subread/1.6.2) was used to count the reads mapping to each gene [52]. The FPKM values for the relative expression of each gene were used for calculating the pairwise AD case/control fold change. Differential expression for pairwise subject comparisons was established as a fold change greater than 1.5.

### Whole genome sequencing and variant calling

DNA was isolated using PureLink Genomic DNA Kit (ThermoFisher). Whole genome sequencing was performed using DNBseq next generation sequencing technology. Libraries were sequenced on an Illumina NovaSeq to generate 100-base, paired-end reads. Sequencing reads were aligned, and variant calls variants were called with the Genome Analysis Toolkit (GATK) Unified Genotyper following the GATK Best Practices 3.3 [53–55].

### Regulatory Element Locus Intersection (RELI)

The RELI algorithm estimated the enrichment of specific genomic features within next generation sequencing datasets, as reported previously [21, 56, 57]. In addition to comparing pairs of datasets (e.g., two ChIP-seq peak sets), RELI systematically estimated the significance of intersections of the genomic coordinates of genetic variants and ChIP-seq peaks, as described previously [21]. In this setting, observed intersection counts are compared to a null distribution composed of variant sets chosen to match the disease loci in terms of the allele frequency of the lead variant, the number of variants in the linkage disequilibrium (LD) block, and the LD block structure.

### Identification of allelic ATAC-seq and ChIP-seq reads using MARIO

To identify possible allele-dependent mechanisms in our functional genomics datasets, we applied our MARIO method [21]. In brief, MARIO identifies common genetic variants that are (1) heterozygous in the assayed cell line (using NGS DNA sequencing data) and (2) located within a peak in a given ChIP-seq or ATAC-seq dataset. MARIO then examines the sequencing reads that map to each heterozygote within each peak for imbalance between the two alleles. We report allelic accessibility and NFKB1 binding at AD genetic risk variants in our ATAC-seq data with an Allelic Reproducibility Score (ARS) greater than or equal to 0.4 which is considered significantly allelic [21].

### Data access

All raw and processed sequencing data generated in this study have been submitted to the NCBI Gene Expression Omnibus (GEO; https://www.ncbi.nlm.nih.gov/geo/) under accession number GSE184238. The reviewer token is urkzwwuszrsnrol. A UCSC Genome Browser session is available at http://genome.ucsc.edu/s/ledsall/AtopicDermatitis.

## ABBREVIATIONS

(ARS): Allelic Reproducibility Score
(ATAC-seq): Assay for Transposase-Accessible Chromatin sequencing
(AD): Atopic dermatitis
(ChIP-seq): Chromatin immunoprecipitation with sequencing
(CCHMC): Cincinnati Children’s Hospital Medical Center
(DICE): Database of Immune Cell Expression, Expression quantitative trait loci and Epigenomics
(EASI): Eczema Area and Severity Index
(eQTL): Expression quantitative trait loci
(FRiP): Fraction of Reads Inside of Peaks
(FPKM): Fragments Per Kilobase of transcript per Million fragments mapped
(GATK): Genome Analysis Tooklit
(GWAS): Genome Wide Association Study
(IGA): Investigator Global Assessment
(LD): Linkage Disequilibrium
(MARIO)method: Measurement of Allelic Ratios Informatics Operator
(PBMCs): Peripheral blood mononuclear cells
(RELI) method: Regulatory Element Locus Intersection
(RNA-seq): RNA sequencing
(STAR): Spliced Transcripts Alignment to a Reference
(TF): Transcription factor
(TSS): Transcription Start Site

## ACKNOWLEDGEMENTS

We thank the contributions of the following physicians in their support in establishing our research clinic: Juan Pablo Abonia, MD, Jonathan Bernstein, MD, Sheharyar Durrani, MD, Stephanie Ward, MD, Justin Greiwe, MD, Michelle Lierl, MD, Kimberly Risma, MD, PhD.

